# Perceiving less or perceiving unreliably? Disentangling thermosensory sensitivity and precision in the contexts of ageing and neuropathy

**DOI:** 10.1101/2025.09.22.677851

**Authors:** Arthur S. Courtin, Camilla E. Krænge, Alexandra G. Mitchell, Jesper Fischer Ehmsen, Peter K. Brask-Thomsen, Sandra Sif Gylfadottir, Francesca Fardo

## Abstract

Thermal perception is determined not only by sensitivity but also by precision. Yet, the latter is often overlooked in thermosensation and pain research. This study examined how ageing and diabetic polyneuropathy (DPN) affect these parameters and whether assessing both sensitivity and precision can aid in distinguishing patients from healthy controls. Using Bayesian hierarchical models, we estimated psychometric function thresholds (sensitivity) and slopes (precision) for cold detection, warm detection, cold pain, and heat pain stimuli delivered at the volar forearm, in a cross-sectional sample of 75 healthy adults (aged 21–80) and 33 patients with DPN. We also estimated these parameters separately for each participant and used the resulting estimates in classification analyses. Ageing was associated with elevated cold and warm detection thresholds, elevated cold pain thresholds, and reduced cold detection slope. Patients with DPN showed similar patterns: higher detection thresholds and lower cold detection slopes while pain-related parameters were largely unaffected. These findings indicate that ageing and neuropathy produce qualitatively similar changes in thermosensory function, particularly affecting cold detection. Classification based on single parameters successfully discriminated patients from controls, except when warm detection or heat pain slopes were used. Combining threshold and slope parameters for a given modality did not significantly improve classification accuracy but combining all parameters across all modalities led to the best performance, with excellent accuracy (AUROCC: .84, 95% CI [.75,.91]). Modelling both thresholds and slopes provides a more comprehensive view of sensory decline and may enhance the detection of early or subtle sensory dysfunction.

**Perspective:** This study shows that ageing and diabetic polyneuropathy produce strikingly similar thermosensory changes, affecting cold detection sensitivity and precision as well as warm detection sensitivity while largely sparing pain-related measures.

## 1. Introduction

Sensitivity and precision are fundamental parameters of sensory processing, determining which stimuli we are able to feel and how we perceive them. Sensitivity can be quantified by psychophysical thresholds, the smallest stimulus intensity that is perceived at least half of the time. Precision, indexed by the slope of the psychometric function relating stimulus intensity to perception probability, reflects the reliability of perceptual judgements ^1–3^. A steep slope indicates a clear boundary between perceived and non-perceived intensities, whereas a shallow slope reflects greater variability in perception. Thresholds are routinely investigated in thermosensation and pain studies but slopes are usually overlooked. However, the assessment of both threshold (sensitivity) and slope (precision) could offer complementary information on how individuals detect and interpret thermal stimuli, potentially revealing differences that remain hidden when focusing on thresholds alone.

Ageing leads to progressive changes in somatosensory function, including thermosensation and nociception. Older adults often exhibit elevated thermal detection and pain thresholds, suggesting that they are less sensitive to temperature changes ^4–7^. These age-related differences may stem from factors such as diminished peripheral nerve conduction velocity, reduced density of cutaneous receptors, and alterations in central sensory processing^8–11^.

Thermal thresholds are often further elevated in patients with diabetic polyneuropathy (DPN), a frequent complication of *diabetes mellitus*^12^. In these patients, chronic hyperglycemia and hyperlipidemia lead to progressive nerve damage affecting all types of peripheral nerve fibres, including those subserving thermosensation and nociception ^13,14^. As a result, individuals with DPN exhibit decreased conduction velocities, reduced intra-epidermal nerve fibre density, and aberrant patterns of spontaneous activity, which contribute to these changes in thermal perception ^12–14^.

Although the literature extensively documents how both healthy ageing and DPN affect thermal thresholds, little research has examined the influence of these conditions on the precision of thermosensory processing. To address this gap, we evaluated the thresholds and slope for cold and warm detection, as well as cold and heat pain perception, across the adult lifespan (20 to 80 years) and between healthy controls and individuals with DPN. We hypothesized that ageing and neuropathy are associated with reduced sensitivity (elevated thresholds) and precision (decreased slope), to different extents for different modalities. Further, we expected that incorporating both measures of perceptual sensitivity and precision may enhance our ability to distinguish patients from healthy controls, ultimately improving assessments of age- and neuropathy-related changes in thermosensory function. These analyses also allowed us to examine whether thermosensory alterations associated with age and neuropathy reflect a shared phenotype or are qualitatively distinct changes, as well as to explore how thresholds and slopes relate to each other across modalities.

## 2. Methods

### 2.1. Design

The study employed a single-session, cross-sectional design to examine the impact of ageing and neuropathy on thermosensation. Participants completed three substudies aimed at assessing thermal sensitivity and precision (detection and pain thresholding), paradoxical heat sensation, and spatial aspects of thermal perception ^15^. In addition, all participants completed survey measures related to handedness ^16^, mental health ^17,18^, and pain sensitivity ^19,20^. Patients also completed surveys on neuropathic pain symptoms ^21,22^.

Each participant’s session lasted approximately four hours and data were collected between April and October 2024. This project was approved by the Region Midtjylland Ethics Committee (Denmark) and adhered to the current version of the Helsinki Convention (2013). Participants provided informed consent prior to their participation in the study. In accordance with Danish ethical guidelines, healthy controls received financial compensation for their participation. Patients were not compensated directly, but the costs associated with their travel and related arrangements were reimbursed.

This paper focuses on a specific subset of the data, related to thermal detection and pain perception across the lifespan and in individuals with diabetic neuropathy.

Patients or the public were not involved in the design or conduct of this experiment.

### 2.2. Participants

To ensure balanced representation across the adult lifespan, the control group was stratified into four age cohorts (20-35, 36-50, 51-65 and 66-80 years) with the aim of including 15 to 25 individuals in each cohort. Patients with DPN were recruited from a pool of patients diagnosed as having probable neuropathy based on the Toronto consensus grading system, as part of other projects within the framework of the Danish National Type 2 Diabetes (DD2) Cohort ^23–27^.

Inclusion criteria required all participants to be between 20 and 80 years old and fluent in Danish and/or English. Exclusion criteria included pregnancy or lactation, recent jet lag or sleep deprivation, non-diabetes-related skin diseases such as eczema or psoriasis, a history of alcohol or drug abuse, cannabis use within the previous four weeks, and alcohol consumption within the previous 48 hours. For healthy controls, having a physical condition and/or medication that might influence pain perception was also an exclusion criteria.

A total of 86 healthy controls and 34 patients with probable diabetic polyneuropathy (DPN) completed the thresholding tasks. Following a secondary review of participant reports and prior to data analysis, we excluded ten control participants due to potential confounding factors: use of medications potentially affecting temperature perception (n = 7), history of epilepsy (n = 1), diagnosis of Reynaud syndrome (n = 1), and diagnosis of ankylosing spondylitis (n = 1). We also excluded one patient due to a history of thermal injury at the stimulation site, related to their professional occupation. After visualizing the data but before formal analysis, one additional control participant was excluded due to poor data quality (see below). A detailed overview of participant recruitment and exclusions is presented in Fig. 1.

**Figure 1.**
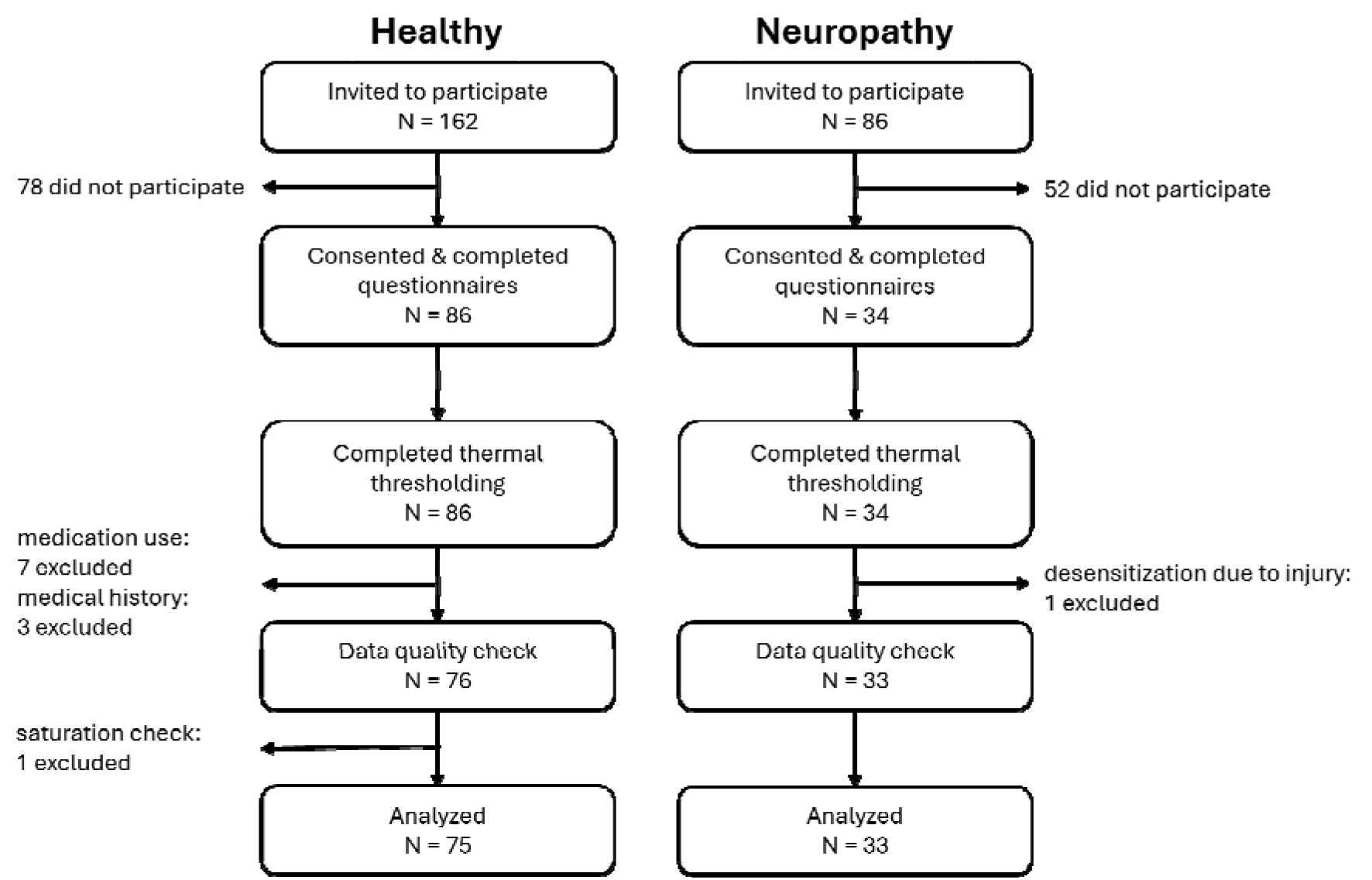
Recruitment flow chart. Recruitment flow chart detailing participant inclusions, exclusions, and final sample sizes for analysis.

The final analyzed sample included 75 healthy controls (30 M and 45 F, self-reported; age range: [21-80]) and 33 patients with neuropathy (18 M and 15 F, self-reported; age range: [47 - 79]; years since diabetes diagnosis as mean±SD: 13.0± 2.2). The proportion of males and females in the different groups is reported in Table S1.

### 2.3. Thresholding tasks

All participants completed four brief computerized tasks designed to assess their ability to detect cooling and warming, as well as their perception of cold and heat pain. We used the data collected during these tasks to quantify the sensitivity and precision of the participant’s thermal perception processes (see below). These tasks were always performed after the end of the surveys and in the same order: cold detection followed by warm detection on the non-dominant volar forearm, then cold pain followed by heat pain on the dominant volar forearm (Fig. 2).

**Figure 2.**
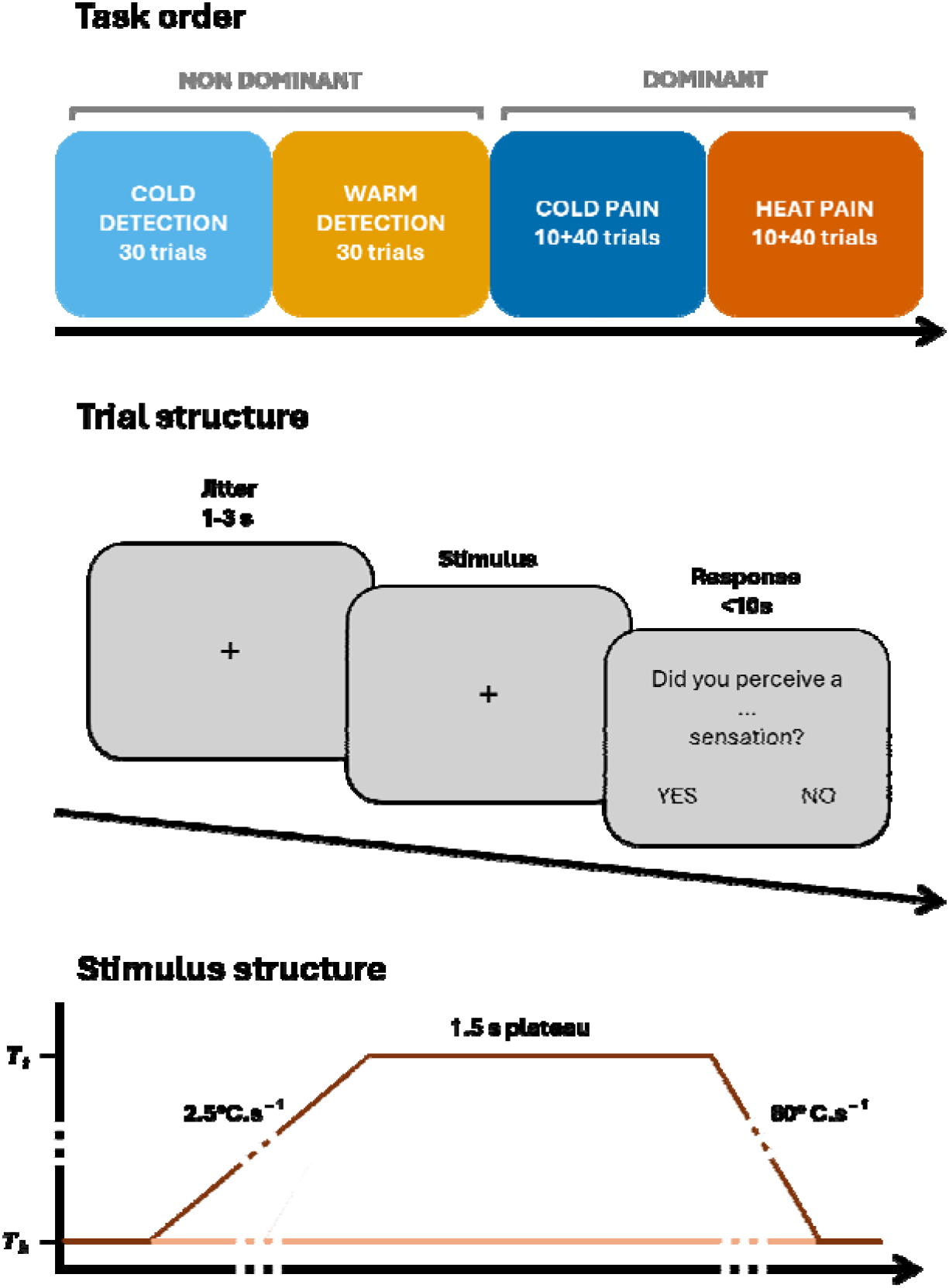
Task design. The top row illustrates the sequence of tasks, including the number of trials for each and the forearm to which the stimulator was applied. The middle row illustrates the structure of a single trial: first, a baseline period, during which the stimulator was held at 30°C for a pseudo-random duration, followed by the stimulus itself, and finally a response period. The bottom row illustrates the temperature profile over time during stimulation: all zones started at baseline temperature T_b_, two zones then reached the target temperature T_t_ at a rate of 2.5°C/s, remained at T_t_ for 1.5 s, and finally returned to T_b_ at a rate of 80°C/s.

All thermal stimuli were delivered using the TCS-II (QST.Lab, Strasbourg, France), a Peltier effect contact stimulator. More specifically, we used the T11 probe which has a total stimulation surface of ∼900 mm^2^ containing five independently controlled zones of 7.4 x 24.2 mm and a maximum rate of temperature change of 100°C/s. For the whole duration of the experiment, all zones of the probe were kept to the neutral temperature of 30°C except during stimulation.

Each trial began with a 1-3 s (randomly sampled) period during which all zones remained at 30°C. This was followed by a change of temperature at a rate of 2.5°C/s in two adjacent stimulation zones. The two contiguous zones were randomly selected from the four possible combinations to ensure that the same skin area was not stimulated in consecutive trials, thus limiting the risk of habituation or sensitization. Once the target temperature was reached (see next section for target temperature definition), it was maintained for 1.5 s, before return to baseline at a rate of 80°C/s. The temperature of each zone was continuously recorded from the onset of the temperature change until the end of the plateau. After each stimulus, participants reported whether they felt cold (yes or no) for cold detection, warm (yes or no) for warm detection, or a burning sensation (yes or no) for pain perception assessment. Their chosen answer was displayed for 0.5 s before the next trial began. For all tasks, participants had a maximum of 10 s to respond, using the left or right arrow keys of the computer’s keyboard (left = yes, right = no).

Each detection thresholding task comprised 30 trials and lasted between 5 and 7 minutes. Each pain thresholding task comprised 10 practice trials followed by 40 test trials and lasted between 10 and 15 minutes. In all cases, the task comprised a mix of trials with infra- and suprathreshold target temperatures (see next section).

For all modalities, participants were informed that each trial would involve a temperature change, followed by a question: “Did you perceive a cold sensation?” for cold detection, “Did you perceive a warm sensation?” for warm detection, and “Did you perceive a burning sensation?” for pain assessments. They were instructed to respond as quickly and accurately as possible. To clarify the concept of ‘burning,’ participants were given examples such as holding a very hot cup of coffee or an ice cube for too long.

In these tasks, participants’ responses summarize their perception during the entire stimulus, including the plateau phase, rather than their instantaneous perception as in the conventional method of limits ^28^. The use of a relatively fast rate of temperature change is therefore unlikely to affect threshold estimates, contrary to what would happen for the conventional method of limits ^29^.

The average ramp duration and proportion of detected stimuli, per participant and task, are reported in Table S2 and S3.

The task was programmed using Psychtoolbox-3 ^30^ in MATLAB R2021a (The MathWorks, USA).

### 2.4. Adaptive stimulus intensity selection

We used an adaptive algorithm to select stimulus intensities on each trial based on the participants’ responses in previous trials of the same task. This approach ensured that stimulus intensities were calibrated to individual sensitivity, limiting the number of trials at uninformative (never or always detected) or excessively painful intensities. As a result, it reduced the number of trials needed to attain a given level of estimation and the risk of habituation or sensitization.

Specifically, we used the Ψ (psi) method, a Bayesian adaptive method of levels thresholding algorithm implemented in Palamedes ^31,32^. This method keeps track of an approximate joint probability distribution for the threshold and the slope of a psychometric function (PF). At the start of each trial, the method uses this probability distribution to select the stimulus intensity that maximizes potential information gain and then updates this distribution based on the participant’s response.

In this case, the method estimated the parameters of a PF relating stimulus temperature to the probability of reporting the perception of interest (cold, warm, cold pain, or heat pain, depending on the task). We modeled the PF as a Gaussian cumulative density function defined by four parameters (threshold, slope, guess rate, lapse rate; Fig. 3). Full implementation details are provided in the Supplementary Materials.

**Figure 3.**
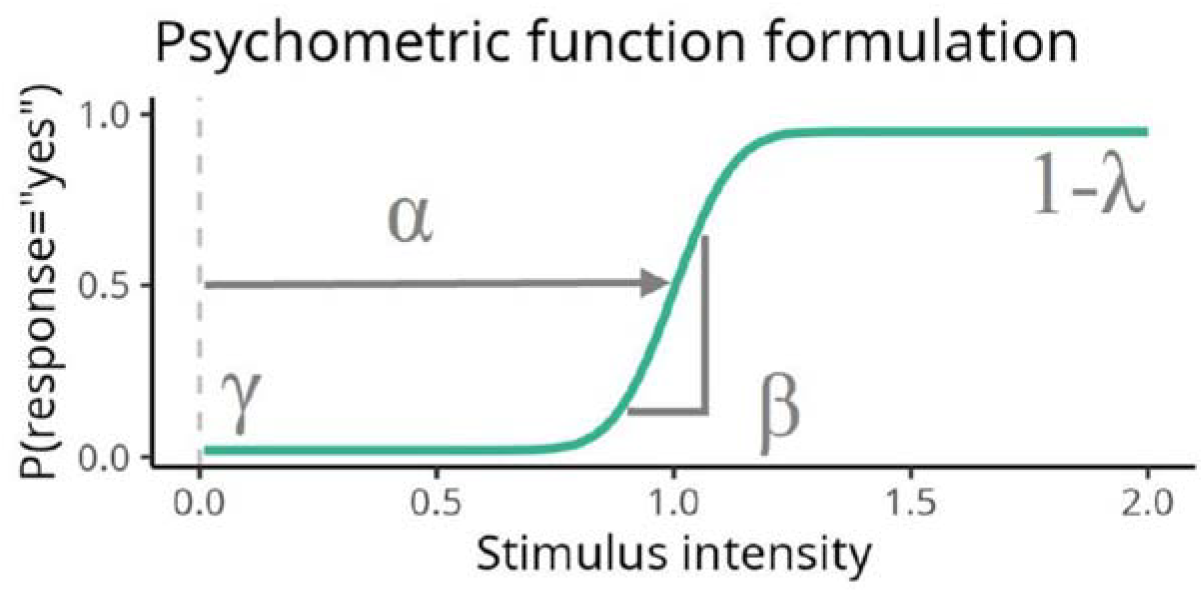
Illustration of the psychometric function formulation. The plot shows how the threshold α, the slope ⍰, the guess rate γ, and the lapse rate λ relate to the psychometric function.

### 2.5. Statistical analyses

The pre-registered data processing and analysis plan for this experiment can be found on the project’s OSF repository: https://osf.io/dvz94.

Deviations from the pre-registered plan were minor and included an adjusted criterion for assessing the quality of temperature recordings, modification of the models due to sampling issues and to address limitations raised during peer-review, and the addition of Bayes Factors to evaluate evidence for the absence of effects in parallel with posterior probability tests. Full details and justifications are available in the OSF repository.

#### 2.5.1. Data quality assessment

We performed several data quality checks before analysis. Neuropathy status was screened using the Michigan Neuropathy Screening Instrument (MNSI) and no control participant exceeded the established cutoff (MNSI≥7) indicative of neuropathy ^19,33^. To account for potential deviations between delivered and target temperatures, we used the probe temperature recorded during the plateau phase of stimulation, rather than the target temperature, as stimulus intensity for data analysis. We excluded trials showing excessive temperature deviations during this period (2 of 15,260 trials deviated by more than 1 °C from the target; 180 of 15,260 trials deviated by more than 0.2 °C from the mean recorded temperature). In control participants, we further verified that detection responses saturated at lower intensities than pain responses; one control participant who violated this criterion was excluded. This criterion was not applied to patients, given the possibility of altered sensory experience due to neuropathy.

#### 2.5.2. Modeling framework

We analyzed the relationship between binary responses (detection or pain) and stimulus temperature using hierarchical Bayesian PF models. These models jointly estimated thresholds (indexing perceptual sensitivity), slopes (indexing perceptual precision), guess rates and lapse rates at the participant and group levels and across thermosensory modalities.

We first examined the effects of age on thermal perception using data from healthy controls, testing whether thresholds increased and slopes decreased with age across modalities while controlling for gender. We then assessed the additional effects of neuropathy by including both patients and controls and modeling participant status together with age and gender. This allowed us to test our hypothesis that neuropathy further increases thresholds and reduces slopes, beyond the effects of ageing alone. Across all analyses, psychometric function specifications were consistent with those used in the adaptive stimulus selection procedure. The models were implemented in Stan and estimated using the *CmdStanR* package for *R* ^34–36^. Full details of model implementation and justification of the chosen priors (Fig S1 ans S2) can be found in the Supplementary Materials.

#### 2.5.3. Interpretation of group- and participant-level parameters

We evaluated preregistered, directional hypotheses using posterior probabilities for group-level effects on psychometric function parameters, with a 0.05 decision threshold analogous to the use of p-values in frequentist inference. Specifically, we tested whether thresholds increased and slopes decreased with age in healthy controls, and whether neuropathy further increased thresholds and reduced slopes beyond the effects of ageing, across thermosensory modalities. We complemented these analyses with Bayes factors for the absence of an effect (BF_01_), computed using the Savage–Dickey density ratio method ^37^ and interpreted using established conventions ^38^. These BF_01_ allow us to distinguish cases where no definitive interpretation can be drawn (no significant difference and inconclusive BF) from cases where there is truly no effect ((no significant difference and sufficient evidence for the absence of an effect).

Beyond group-level differences, we investigated whether individual thresholds and slopes derived from psychometric functions for cold detection, warm detection, cold pain, and heat pain could reliably distinguish patients from healthy controls. Participant-level parameters were estimated separately for each modality and entered into age-adjusted logistic regression models predicting group membership from threshold, slope, or their combination. To avoid biasing the estimation of the classification performance, we only included in this analysis control participants who were at least as old as the youngest patient. Classification performance was quantified using the area under the receiver operating characteristic curve (AUROCC) and tested against chance level. We further assessed whether thresholds and slopes provided complementary information within and across thermosensory modalities.

Finally, to investigate associations between psychometric parameters independent of the effects of gender, age or neuropathy, we inspected the correlation matrix estimated within the hierarchical models. A correlation coefficient was deemed significant when its 95% credible interval did not include 0.

Technical details on the implementation of those tests can be found in Supplementary Materials.

#### 2.5.4. Sample size justifications

The sample size was selected based on previous experiments reported in the literature and available resources. There is no analytical solution for sample size calculation for Bayesian hierarchical models. Simulation-based sample size analyses are possible but computationally expensive and the validity of their results is dependent on the accuracy of arbitrary choices when setting up the simulations ^39^.

## 3. Results

### 3.1. Effects of age on temperature perception

Increasing age was associated with significantly higher cold detection, warm detection and cold pain thresholds (expressed as absolute deviation from baseline skin temperature), indicating an overall reduction in thermal sensitivity (Table 1; Fig. 4). In addition, age was significantly associated with reduced sensory precision, as reflected by a decrease in the slope of the cold detection psychometric function (Table 1). By contrast, effects of age on heat pain thresholds and on slopes for warm detection and pain modalities were inconclusive, providing no clear evidence for systematic age-related changes in these parameters (Table 1). To visualize these effects, Fig. 4 displays group-level psychometric functions estimated at ages 20 and 80, corresponding to the minimum and maximum ages allowed by the inclusion criteria.

**Table 1.** Effects of age on psychometric function parameters across modalities.

| Modality | Parameter | $\kappa$ | 90% CI | $P(H_0)$ | $\log_{10}(BF_{01})$ | Interpretation |
| --- | --- | --- | --- | --- | --- | --- |
| Cold Detection | Threshold | 0.015 | [0.005, 0.025] | .009 | -0.86 | ↗ with age |
|  | Slope | -0.016 | [-0.027, -0.005] | .009 | -0.74 | ↗ with age |
| Warm Detection | Threshold | 0.009 | [0.003, 0.014] | .006 | -0.94 | ↗ with age |
|  | Slope | -0.004 | [-0.013, 0.006] | .242 | 0.25 | Inconclusive |
| Cold Pain | Threshold | 0.006 | [0.001, 0.011] | .016 | -0.68 | ↗ with age |
|  | Slope | 0.000 | [-0.006, 0.006] | .501 | 0.35 | Inconclusive |
| Heat Pain | Threshold | 0.000 | [-0.001, 0.002] | .261 | 0.46 | Inconclusive |
|  | Slope | 0.001 | [-0.009, 0.011] | .577 | 0.37 | Inconclusive |

$\kappa$ denotes the posterior mean of the age effect in log-space. CI = credible interval. $P(H_0)$ denotes the posterior probability that the age effect is equal to 0 or in the direction opposite to the hypothesized effect. $BF_{01}$ is the Bayes factor in favor of the absence of an effect, reported on a $\log_{10}$ scale. Arrows indicate the direction of the age-related effect.

**Figure 4.**
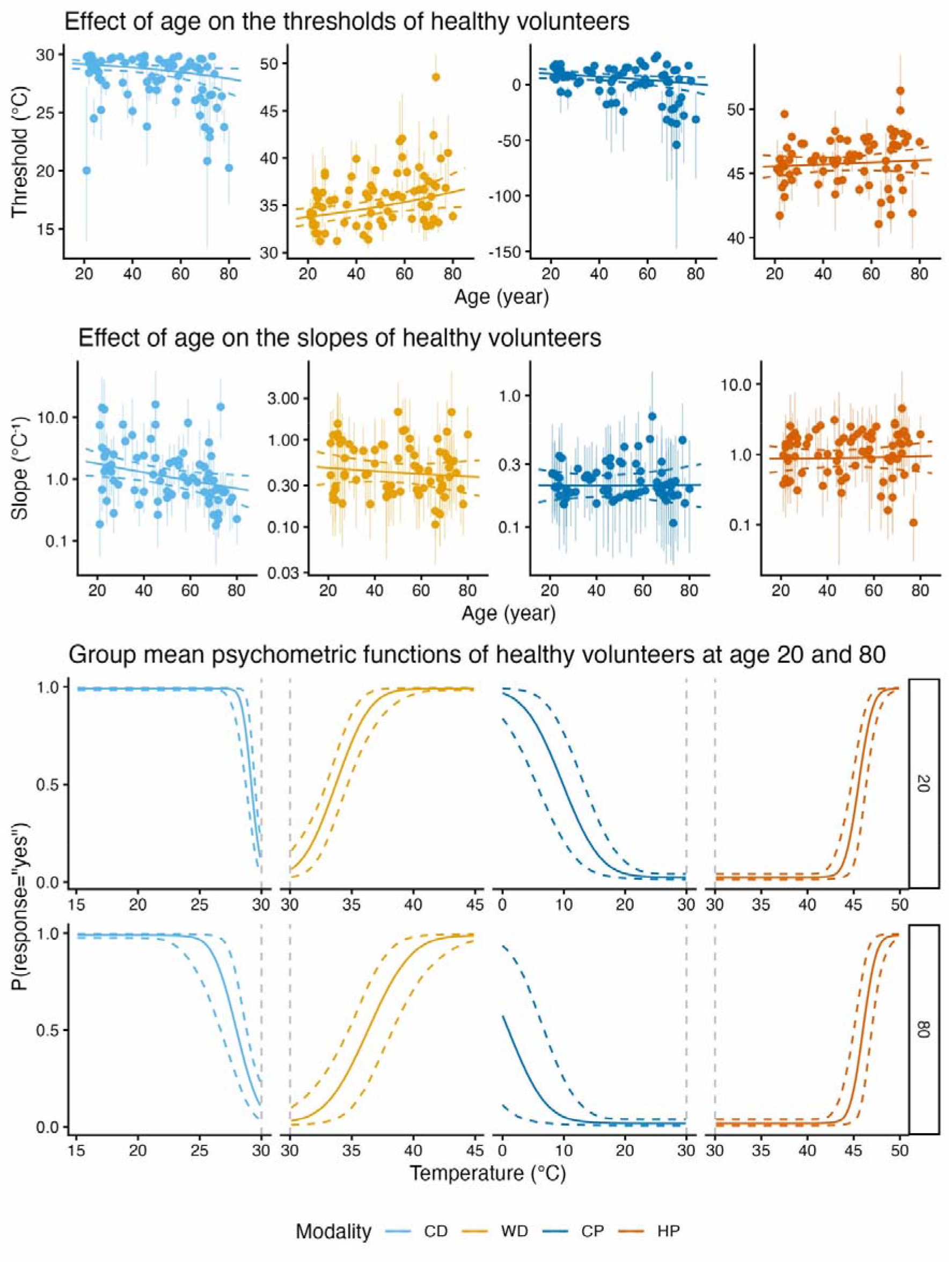
Effect of age on psychometric function parameters. Individual participants’ threshold (top row) and slope (second row) estimates plotted against age, across four sensory modalities: cold detection (CD), warm detection (WD), cold pain (CP), and heat pain (HP). Notice the logarithmic scaling of the y-axis. We also display the group-level psychometric functions estimated at ages 20 and 80 for each modality (bottom rows). Shaded areas represent 95% credible intervals (CI).

### 3.2. Effects of neuropathy on temperature perception

Patients with neuropathy showed significantly higher cold detection and warm detection thresholds compared to age-matched controls, indicating reduced sensitivity to innocuous thermal stimuli (Table 2; Fig. 5). Cold detection slopes were also lower in patients, indicating less reliable perception (Table 2; Fig. 5). In contrast, there was substantial evidence supporting the absence of an effect of neuropathy on heat pain threshold. Comparisons of the cold pain thresholds, as well as slopes for warm detection, cold pain, and heat pain between patients and controls were inconclusive. To illustrate these effects, group-level psychometric functions were estimated at age 50 for patients and controls (Fig. 5). Representations of individual psychometric functions plotted against the participant’s data are available in the Supplementary Materials (Fig S3 to S6).

**Table 2.**
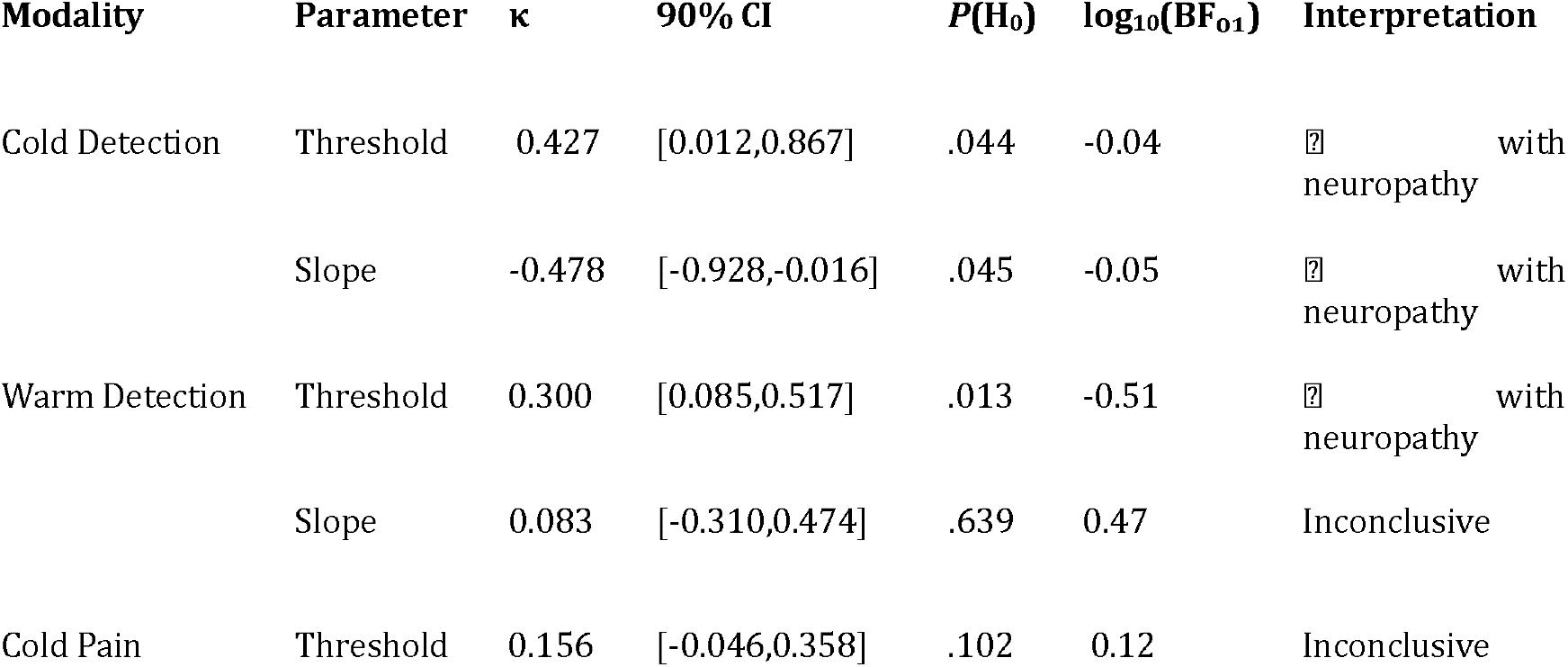
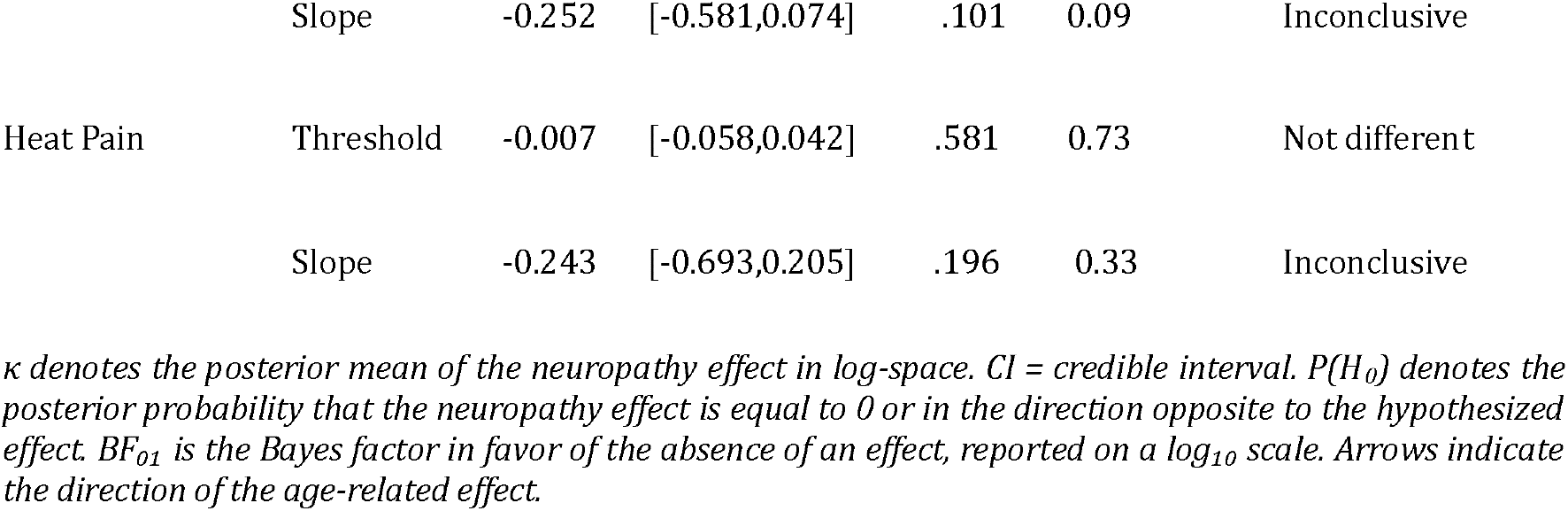
Effects of neuropathy on psychometric function parameters across modalities.

**Figure 5.**
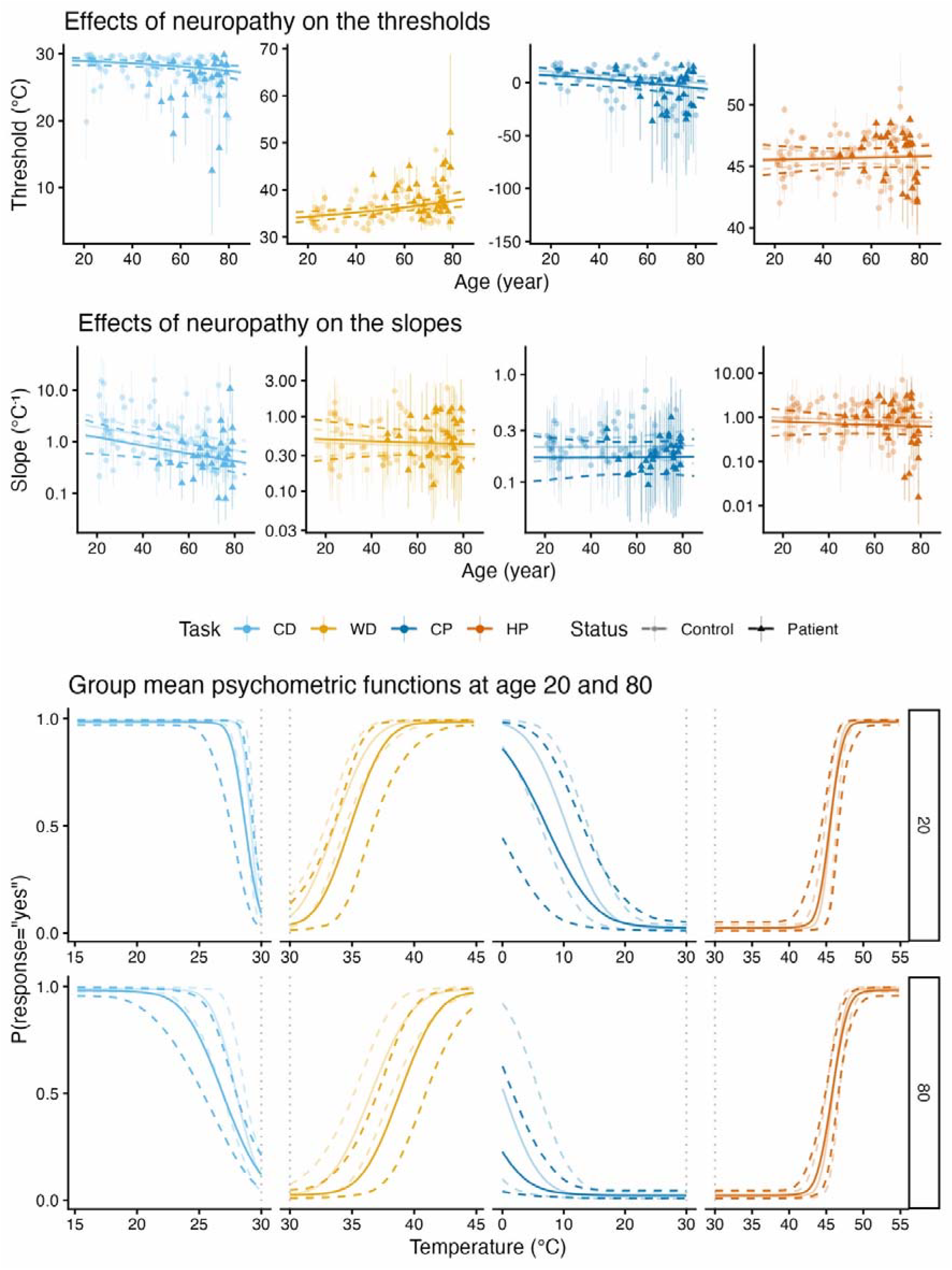
Effect of neuropathy on psychometric function parameters. Individual threshold (top row) and slope (second row) estimates for healthy controls (light) and patients (dark) plotted against age, across four sensory modalities: cold detection (CD), warm detection (WD), cold pain (CP), and heat pain (HP). Notice the logarithmic scaling of the y-axis. We also display the group-level psychometric functions estimated at ages 20 and 80 for each modality (bottom rows). Shaded areas represent 95% credible intervals (CI).

### 3.3. Effects of gender on temperature perception

We never observed a significant difference between males and females, with a number of tests showing substantial evidence for the absence of an effect (Table S4 and S5).

### 3.4. Discrimination between healthy controls and patients with neuropathy

PF parameters afforded varying levels of discrimination between patients and controls of similar age, with AUROCC values ranging from .47 to .84, depending on the modality and whether threshold, slope, or both were included in the score used for classification (Fig. 6, Fig. S7 and S8). Classification performance exceeded chance level, as indicated 90% by confidence intervals that did not include .5, for all tested parameter combinations except for models containing only warm detection slope or heat pain slope.

**Figure 6.**
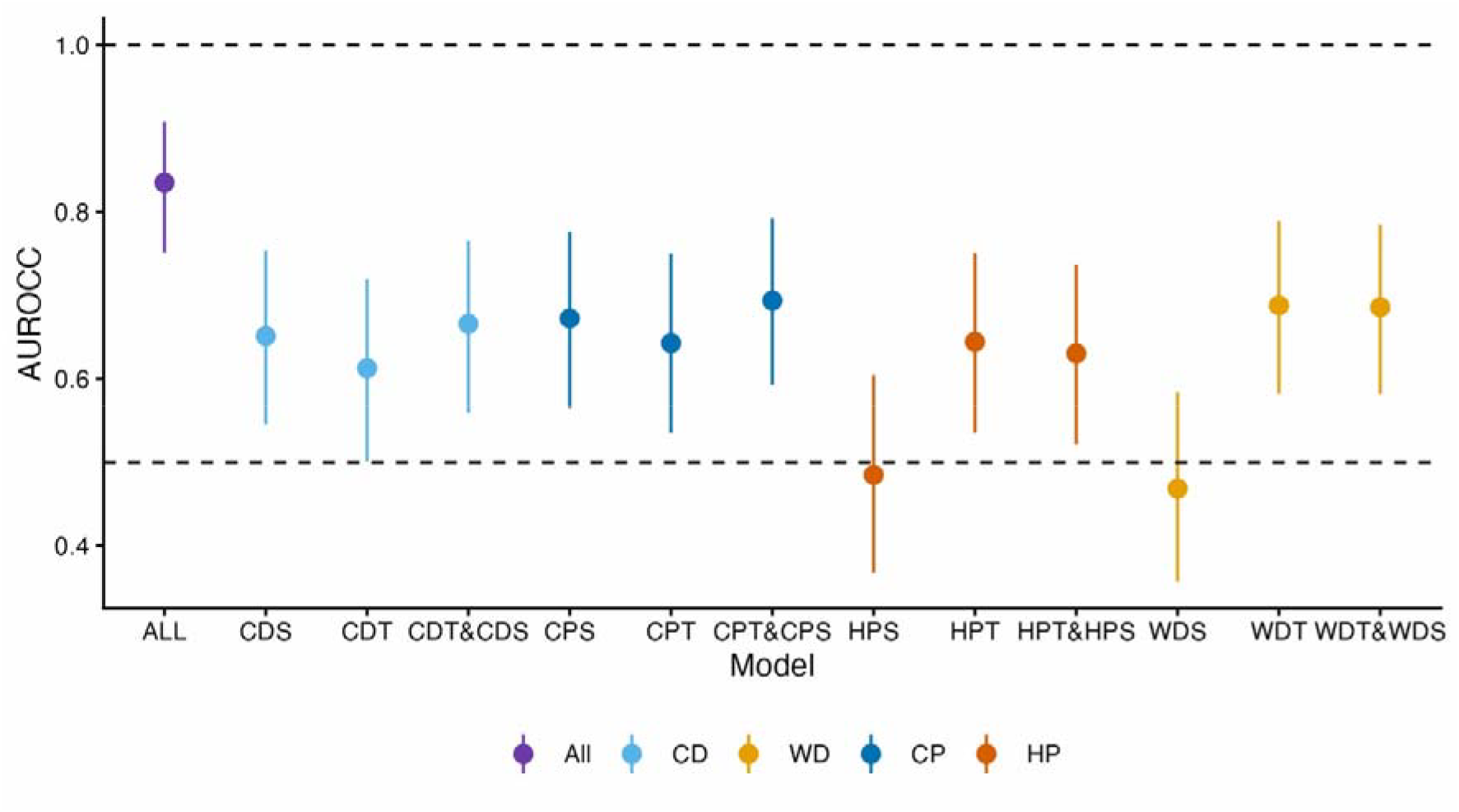
Discriminability of patients and controls based on combinations of psychometric function parameters. The figure shows the observed AUROCC (dots) and their 90% CI (line range) for a variety of scores including only the slope (S), only the threshold (T), or a combination of the threshold and slope for each modality: cold detection (CD), warm detection (WD), cold pain (CP), and heat pain (HP). We also display the AUROCC and 90% CI for a score including thresholds and slopes for all modalities. Scores were corrected for age.

Within each modality, classification performance appeared to reach a plateau once the most informative parameter was included in the score. For warm detection, classification based on the combined threshold and slope score outperformed classification based on slope alone (D = -2.00, p = .023) but not classification based on threshold alone (D = 0.13, p = .552). For the other tasks, classification accuracy using the combined threshold and slope score was not significantly better than classification based on threshold alone (cold detection: D = -0.93, p = .176; cold pain: D = -1.21, p = .114; heat pain: D = 0.84, p = .800) or slope alone (cold detection: D = -0.37, p = .356; cold pain: D = -0.67, p = .251; heat pain: D = -1.31, p = .095).

### 3.5. Association between parameters

Partial correlation analysis, controlled for age and patient status, revealed significant negative correlations between cold detection thresholds and slopes and between cold pain thresholds and slopes, as well as positive correlations between cold and warm detection thresholds, cold and heat pain thresholds, and heat pain thresholds and slopes (Fig. 7). All other correlations did not significantly differ from 0.

**Figure 7.**
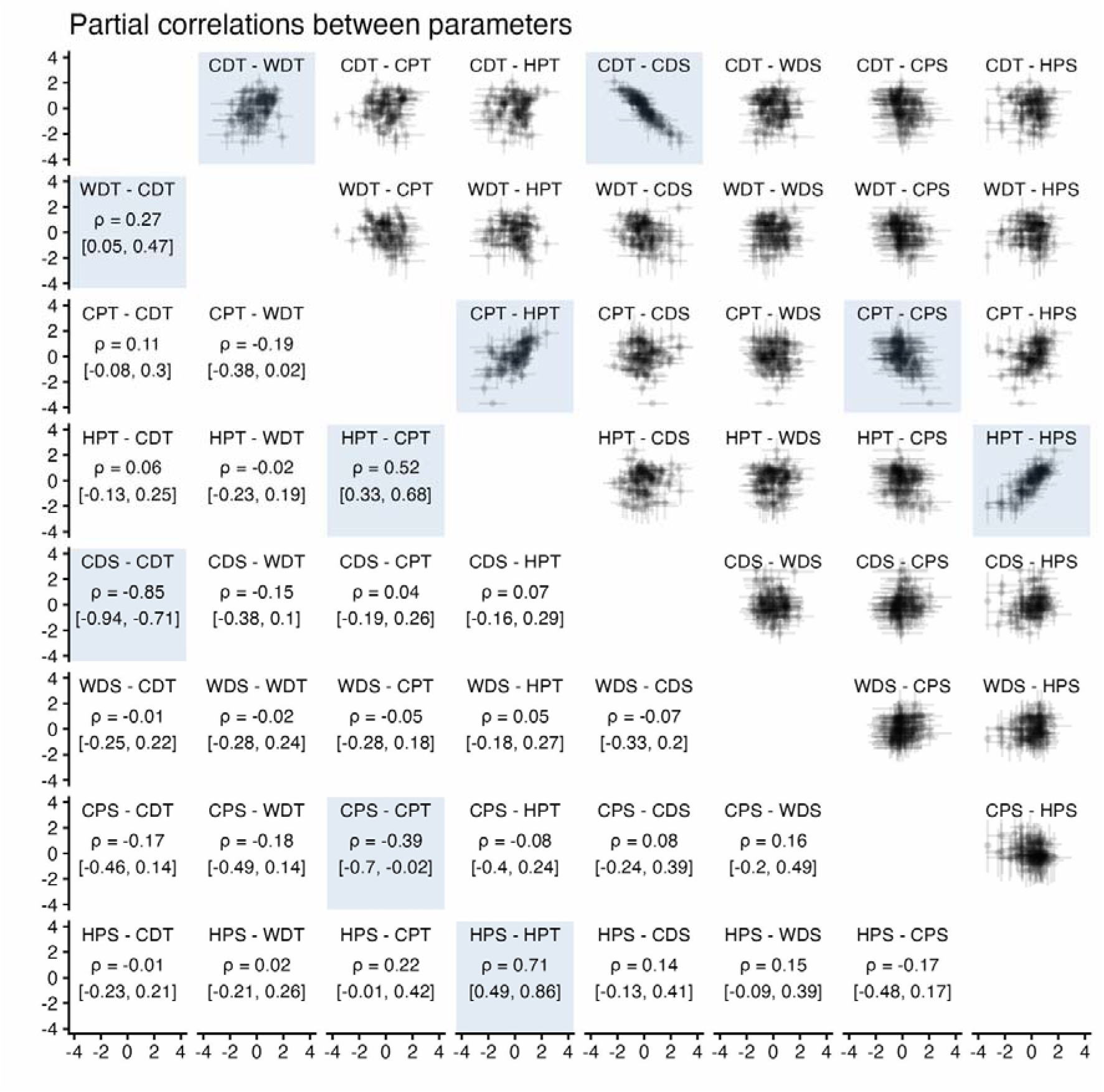
Partial correlation matrix. The upper triangle of the matrix displays scatter plots of standardised residual estimates for each participant and pair of parameters. The dots correspond to posterior means and the lines to 95% CIs. The lower triangle displays correlation coefficients posterior means (ρ rho) and 95% CIs.

## 4. Discussion

This study investigated how ageing and diabetic polyneuropathy (DPN) affect the sensitivity and precision of thermal perception, using the thresholds and slopes of psychometric functions across four modalities: cold detection, warm detection, cold pain, and heat pain. We also examined whether these parameters could differentiate patients with DPN from healthy controls and whether they correlated with each other.

We report four key findings. First, ageing was associated with higher thresholds in multiple modalities and flatter cold detection slopes. Second, DPN had a similar effect, increasing detection thresholds and flattening cold detection slopes. Third, accurate discrimination between patients and controls was possible based on most PF parameters, with a score combining all parameters achieving excellent classification. Finally, detection and pain parameters appeared to form two separate ensembles, with significant correlations within but not between them.

Consistent with previous findings, we observed age-related increases in thermal detection and cold pain thresholds, suggesting reduced sensitivity to temperature changes with increasing age ^4^. Importantly, our results extend these findings by showing that ageing reduces the precision or reliability of cold detection, as indicated by a flatter psychometric function (reduced slope). This means that older adults not only require more intense stimuli to detect skin cooling but also experience greater variability in whether and how they perceive it.

We found no significant age-related effects on the slope of warm detection or pain modalities, as well as on heat pain thresholds. This may suggest that ageing disproportionately affects specific thermosensory pathways, potentially due to differences in small-fiber subtype vulnerability or central processing. However, the Bayes Factors indicated insufficient evidence to conclude a true absence of effect, so these null findings may also reflect limited power due to greater interindividual variability.

On average, patients with DPN had higher cold and warm detection thresholds and lower cold detection slopes compared to age-matched controls. These results are consistent with known peripheral nerve fiber degeneration in DPN ^40–42^. Interestingly, pain thresholds and slopes were either clearly not affected by neuropathy (heat pain threshold) or not clearly different between patients and controls (cold pain threshold and slope, heat pain slope). One possibility is that nociceptive pathways are less susceptible to neuropathic damage or more variable in their impairment, reducing our statistical power to detect these changes. Alternatively, heat pain thresholds may have been truly unaffected in our sample, contrasting with a large number of studies finding heat pain threshold increases in DPN ^43^. This discrepancy could reflect differences in the test site, as DPN is thought to be length-dependent and we tested the volar forearm, a more proximal location than the foot dorsum used in most prior studies ^13^. Another explanation could come from methodological differences, as previous studies used the conventional method of limits to measure the thresholds. In such tasks, a continuously increasing stimulus is applied and participants press a button as soon as they perceive it ^28^. The average intensity at the time of button press across three trials is taken as the threshold measure. Because this depends on response timing, differences in response speed can bias the threshold estimate, even when perception is identical. Moreover, since perception can occur well above or below the actual threshold and since only a few trials are averaged, the estimate is variably influenced by both the true threshold and the slope. In contrast, our tasks do not rely on when participants reply, preventing contamination by response time, while jointly estimating thresholds and slopes, preventing their conflation. Eliminating these confounds may explain why we found substantial evidence against an effect of neuropathy on heat pain thresholds.

Taken together, our results suggest that the effects of DPN on thermal detection closely parallel those observed in healthy ageing, both in direction and magnitude. For cold detection thresholds, cold detection slopes and warm detection thresholds (the three parameters which were significantly affected by neuropathy) the impact of DPN was roughly equivalent to the impact of 30 years of healthy ageing. This alignment suggests that neuropathy may accelerate ageing-like changes in the thermosensory system, rather than induce a qualitatively different pattern of impairment. In this view, DPN compresses or amplifies the trajectory of sensory decline typically seen with age. This interpretation aligns with broader theories proposing that chronic conditions such as diabetes mimic or hasten normal ageing processes in the peripheral and central nervous systems ^44,45^. However, because our study is cross-sectional, we cannot determine whether neuropathic alterations truly reflect accelerated ageing in thermal perception. Longitudinal data will be needed to test this hypothesis directly.

Discrimination performance was primarily driven by a subset of PF parameters. Classification improved further only when parameters were combined across modalities, whereas combining thresholds and slopes within a single modality did not significantly enhance performance when either of them led to better than chance classification. This pattern demonstrates the added value of combining information across modalities in distinguishing patients from controls. The AUROCC values we obtained for classification based on cold (.61 [.50,.72] vs .69 [.63,.75]) or warm detection thresholds (.69 [.75,.79] vs .66 [.60,.72]) alone were in line with those reported in a previous study using conventional method of limits measurements ^46^.

Finally, beyond classification performance, we examined associations between thermal thresholds and PF slopes. While many studies have investigated relationships between thermal thresholds and biological or psychological variables (e.g., ^47–50^), few have focused on how different thermal thresholds relate to one another, or to PF slopes, in the absence of external explanatory factors. Our partial correlation analysis revealed several associations among these parameters, beyond shared effects of neuropathy or age. Interestingly, significant correlations were only observed between detection metrics or between pain metrics. This indicates a certain separation between pure thermosensation and thermonociception, rather than a smooth continuum encompassing all temperature perception. These findings also suggest that assessing both cold and warm detection (or pain) parameters does not provide entirely independent information, which could have implications for test selection in clinical or experimental settings.

Taken together, our results suggest that replacing the conventional method of limits with an adaptive method of levels in QST protocols may be advantageous, as it likely enables more accurate estimation of thresholds and allows estimation of psychometric function slopes. However, clinical adoption depends not only on potential benefit but also on the cost-benefit ratio. In this respect, the relatively long testing duration of approximately 5 to 7 minutes for detection and 10 to 15 minutes for pain may represent a high cost relative to the potential gain. Future studies should examine whether optimized versions of the adaptive method can achieve comparable or superior estimation precision with fewer trials and therefore shorter testing times. In the current implementation, the adaptive method was initialized with flat priors over a wide range of parameter values. Estimation efficiency could likely be substantially improved by constraining the parameter space and incorporating informative priors derived from previous datasets, such as the present one.

## 5. Conclusion

In summary, our results indicate that ageing and diabetic polyneuropathy weaken thermosensory function in strikingly similar ways: both elevate cold and warm detection thresholds and flatten the cold detection slope, while leaving pain measures largely intact. This overlap suggests that neuropathy accelerates the normal trajectory of sensory ageing rather than introducing a fundamentally different pattern of impairment. By modelling thresholds and slopes together, our study provides a more comprehensive picture of thermosensory decline and lays the groundwork for refined quantitative sensory tests that can better track neuropathic changes.

## Acknowledgments

The authors would like to thank Johanne Sejrskild Rejsenhus for her help with data collection.

## Disclosure

### Author contributions

Conceptualization: ASC, AGM, CEK, FF

Data curation: ASC, CEK, AGM

Formal analysis: ASC

Funding acquisition: FF

Investigation: AGM, CEK

Methodology: ASC, AGM, CEK, JFE, FF

Project administration: CEK, FF

Resources: PBT, SSG

Software: ASC, AGM

Supervision: FF

Visualization: ASC

Writing – original draft: ASC, FF

Writing – review & editing: ASC, AGM, CEK, JFE, FF, SSG, PBT

### Data and materials availability

Data and analysis code for this study is available at

https://github.com/Body-Pain-Perception-Lab/lifespan-and-neuropathy-sensitivity-and-precision

### Responsible use of AI

During the preparation of this work the authors used ChatGPT to edit and improve the text of the manuscript. After using this service, the authors reviewed and further edited the content. They take full responsibility for the content of the published article

