## supplementary materials for "Perceiving less or perceiving unreliably? Disentangling thermosensory sensitivity and precision in the contexts of ageing and neuropathy"

Adaptive Method and Data Analysis Implementation

### Bayesian Ψ (Psi) adaptive method implementation

In all cases, the algorithm modeled the PF as a Gaussian cumulative density function:

$$\Theta(x;\alpha,\beta,\gamma,\lambda)=\gamma+(1-\gamma-\lambda) \Phi\left( \beta(x-\alpha) \right)$$

*where Θ is the probability of reporting the target perception, x is the absolute temperature deviation from baseline, α is the threshold, β is the slope, λ is the lapse rate, γ is the guess rate, and Φ is the standard Gaussian distribution cumulative density function*

The guess and lapse rates, corresponding to the probability of reporting a detection at very low intensities and not reporting a detection at very high intensities respectively, were both set to .05. For cold detection and cold pain assessments, possible stimulus values ranged from 29.9℃ to 0.1℃ in 0.1℃ increments. For warm detection and heat pain assessments, possible stimulus intensities ranged from 30.1℃ to 50℃ in 0.1℃ increments. In all cases, the algorithm kept track of the approximate joint posterior distribution for slope and threshold values on a 51 by 51 grid, with possible threshold values uniformly distributed across the stimulus range and possible $log_{10}$slope values uniformly distributed between $log_{10}(0.001^{\circ}C^{-1})$and $log_{10}(6^{\circ}C^{-1})$(inclusive). We used uniform priors for all parameters. Out of the 50 trials used to assess pain sensitivity, the first 10 were familiarization trials, for which stimulus intensities were selected using an up-down staircase algorithm. This ensured that the participant had some time to stabilize their strategy regarding what they reported as burning. These trials were not included in the analysis.

### Model implementation

The dataset consisted of stimulus temperatures and the binary yes/no responses provided by participants, with responses grouped by modality within participants. To estimate thresholds and slopes for each modality and participant while accounting for this nested structure, we used hierarchical Bayesian psychometric function (PF) models.

#### Effects of age on thermal perception

We first assessed how ageing affects thermal perception, limiting our analyses to data from healthy controls. Specifically, we tested whether thresholds increase and slopes decrease with age, across thermosensory modalities.

We assumed a Gaussian PF, in line with the parametrization of the adaptive method. Thresholds and slopes were modelled as varying across participants and modalities. Guess and lapse rates were allowed to vary between participants but remained constant across modalities, as they represent an individual’s overall tendency to make errors. All PF parameters were modeled as affine functions of the participant’s age and gender, with a random intercept. To improve model interpretability, age was centered at 20 years, ensuring that the intercept corresponded to parameter values for the youngest possible participant. Participant gender was indexed by a dummy variable (-0.5 for females, 0.5 for males). Thresholds and slopes were modelled in log-space to ensure that they were positive, reflecting the assumptions that an increase in stimulus intensity is associated with an increase of the probability of perception. Guess and lapse rates were modelled in logit space, constraining their values to the 0 to 0.5 range.

$$\Theta(x;\alpha,\beta,\gamma,\lambda)=\gamma+(1-\gamma-\lambda) \Phi\left( \beta(x-\alpha) \right)$$

$$ln(\alpha)=\kappa_{0,\alpha}+\kappa_{a,\alpha} Age +\kappa_{g,\alpha}Gender+\epsilon_{i,\alpha}$$

$$ln(\beta)=\kappa_{0,\beta} +\kappa_{a,\beta}Age +\kappa_{g,\beta}Gender+\epsilon_{i,\beta}$$

$$logit(2\gamma)=\kappa_{0,\gamma} +\kappa_{a,\gamma}Age+\kappa_{g,\gamma}Gender+\epsilon_{i,\gamma}$$

$$logit(2\lambda)=\kappa_{0,\lambda} +\kappa_{a,\lambda}Age+\kappa_{g,\lambda}Gender+\epsilon_{i,\lambda}$$

*where Θ is the probability of reporting the target perception, x is the absolute temperature deviation from baseline, α is the threshold, β is the slope, λ is the lapse rate, γ is the guess rate, Φ is the standard Gaussian distribution cumulative density function, 𝜅’s are main effects and ϵ’s are random intercept effects for participant i.*

#### Effects of neuropathy on thermal perception

After assessing the effects of ageing in healthy controls, we investigated how neuropathy alters thermal perception by including data from both healthy controls and patients. This allowed us to test our hypothesis that neuropathy further increases thresholds and reduces slopes, beyond the effects of ageing or gender alone.

To account for the combined effects of age, gender, and neuropathy, the models described in the previous section were extended to differentiate between healthy controls and patients. Participant status was indexed by a dummy variable (-0.5 for controls, 0.5 for patients).

$$ln(\alpha)=\kappa_{0,\alpha}+\kappa_{s,\alpha} Status+\kappa_{a,\alpha} Age +\kappa_{g,\alpha}Gender+\epsilon_{i,\alpha}$$

$$ln(\beta)=\kappa_{0,\beta} +\kappa_{s,\beta} Status+\kappa_{a,\beta}Age +\kappa_{g,\beta}Gender+\epsilon_{i,\beta}$$

$$logit(2\gamma)=\kappa_{0,\gamma} +\kappa_{s,\gamma} Status+\kappa_{a,\gamma}Age+\kappa_{g,\gamma}Gender+\epsilon_{i,\gamma}$$

$$logit(2\lambda)=\kappa_{0,\lambda} +\kappa_{s,\lambda} Status+\kappa_{a,\lambda}Age+\kappa_{g,\lambda}Gender+\epsilon_{i,\lambda}$$

#### Model estimation

The models were implemented in *Stan* and estimated using the *CmdStanR* package for *R*, using the *Rstudio* interface^1–3^. Each model was sampled using four chains of 2,000 warm-up and 2,000 sampling iterations, with a target average acceptance probability of .95 and a maximum tree depth of 12. Initial values for Stan were generated from a quick *pathfinder* optimization using default parameters except for the fact that resampling was turned off.

To ensure proper model sampling, we relied on a series of conventional diagnostic checks ^3^. None of the post-warm-up chains showed divergent transitions and all Estimated Bayesian Fractions of Missing Information (E-BFMI) were higher than 0.3, indicating adequate sampling of the posterior. All parameters had $\hat{R}$ diagnostic values below the conventional cut-off of 1.05, indicating convergence of the chains to a common solution. Finally, all group-level parameters had sufficient bulk and tail effective sample sizes (bulk-ESS and tail-ESS), indicating an adequate number of independent draws for inference.

### Interpretation of group-level parameters

#### Hypothesis tests using posterior probabilities

Pre-registered hypothesis tests were conducted by comparing the posterior probability of null hypotheses (H0) to a .05 cut-off. This inference strategy is similar to the use of p-values in frequentist statistics.

To test the prediction that thresholds increase with age, we assessed whether the group-level age effects on thresholds ($\kappa_{a,\alpha}$ for cold detection, warm detection, cold pain, and heat pain) were smaller than or equal to zero, using estimates from the ageing only models.

To test whether the slope of thermal perception decreases with age, we assessed whether the group-level age effect on slopes in the ageing only models ($\kappa_{a,\beta}$ for all modalities) were larger than or equal to zero.

To test the prediction that patients with neuropathy have higher thresholds than age-matched healthy controls, we assessed whether the group-level effects of neuropathy on thresholds ($\kappa_{s,\alpha}$ for all modalities) were smaller than or equal to zero, using estimates from the ageing and neuropathy models.

Finally, to test whether neuropathy increases the variability of thermal perception, we assessed whether the group-level effects of neuropathy on slopes ($\kappa_{s,\beta}$ for all modalities) were larger than or equal to zero, using estimates from the ageing and neuropathy models.

#### Tests for an absence of effect using Bayes Factors

As some of the tests described in the previous section were not significant, we also computed Bayes factors for the absence of an effect (i.e., BF_01_) using the Savage-Dickey density ratio method ^4^. Following Jeffrey’s scale, we interpreted log_10_(BF_01_) values between -0.5 and 0.5 as indicating insufficient evidence to draw conclusions. Values above 0.5 were taken as substantial evidence for the absence of an effect, while values below -0.5 were interpreted as substantial evidence for the presence of an effect ^5^.

### Analysis of participant-level parameters

#### Discrimination between healthy controls and patients

We investigated whether the thresholds and slopes of the psychometric functions (PFs) for cold detection, warm detection, cold pain, and heat pain provided complementary information for distinguishing patients from healthy controls, beyond group-level differences in these parameters. Classification accuracy was evaluated using age-adjusted scores derived from thresholds, slopes, or their combination, either within individual modalities or across all modalities, and tested against chance performance (.5).

PF parameters were estimated independently for each participant and modality using the same estimation procedure as for the hierarchical group models, except that the target average acceptance probability was set to .99. This approach approximates a clinical scenario in which a physician has access only to an individual patient’s data and compares it against normative cut-off values. Individual parameter estimates were entered into logistic regression models (*glm* in R) predicting group membership from threshold and/or slope, while controlling for age. Classification performance was quantified using the Area Under the Receiver Operating Characteristic Curve (AUROCC), computed with the *pROC* package. AUROCC values were tested against .5 by computing their 90% CI (one-sided test) and checking whether it included that value. For each modality, models including both threshold and slope were compared with models including only one parameter to assess whether thresholds and slopes contributed independent information. All comparisons were conducted using *roc.test* with 10,000 stratified resampling iterations. We only included control participants who were at least as old as the youngest patient in this analysis, to avoid biasing the discrmination accuracy estimates due to many easy to classify young controls.

#### Association between PF parameters

To investigate associations between PF parameters beyond the effects of age and neuropathy, we inspected the correlation matrix estimated within the hierarchical model. This reflects the associations between parameters after accounting for these explanatory variables. We summarized the posterior draws for each cell of the residual correlation matrix as the mean and the .025 and .975 quantiles, providing estimates of the correlation coefficients and their 95% CI.

Justification of the priors

Below, we outline the principles used to define the priors used in our models.

### Ageing only model

The priors for the intercepts (i.e., values at age 20) were informed by a previous dataset of 43 healthy participants aged 18–39, collected using closely matched procedures. This dataset was first fitted using a mode structurally identical to those used in the present study, but excluding age effects (i.e., no age-related coefficients).

For this initial fit, we used weakly informative priors for the group-level means, with prior mass concentrated in plausible ranges: 0–10℃ for detection thresholds, 10–20℃ for pain thresholds, values corresponding to psychometric slopes spanning <1℃ to >30℃, and guess/lapse rates between 0.1% and 5%. Between-subject standard deviations were given half-normal priors with mean 0 and the same SD as the corresponding group-level prior.

Posterior distributions from this initial fit were then used to define the priors for the models in this paper. For intercepts of key parameters (thresholds and slopes), we diluted the influence of prior data by inflating the posterior SD (analogous to a SEM) by a factor of four. For nuisance parameters (guess and lapse rates), we used the undiluted posterior distributions directly, as they closely matched the original weakly informative priors. The psychometric function entailed by these priors are represented in Fig. S1.

The priors on between-subject standard deviations in the final models followed the same principles as for group-level means: they were specified as half-normal distributions, but centered on the posterior mean SDs obtained from the earlier fit, with the scale (SD) inflated by a factor of four for threshold and slope parameters, and left unchanged for guess and lapse rates.

Priors for age effects on psychometric parameters had a mean of zero and a standard deviation equal to one sixtyth of the posterior mean for the corresponding between-subject SD in the initial fit. This is roughly equivalent to setting a standard Gaussian prior on the standardized mean difference (Cohen’s *d*) between age 20 and 80. Similarly, priors for gender effects on psychometric parameters had a mean of zero and a standard deviation equal to the posterior mean for the corresponding between-subject SD in the initial fit. This is roughly equivalent to setting a standard Gaussian prior on the standardized mean difference (Cohen’s *d*) between males and females.

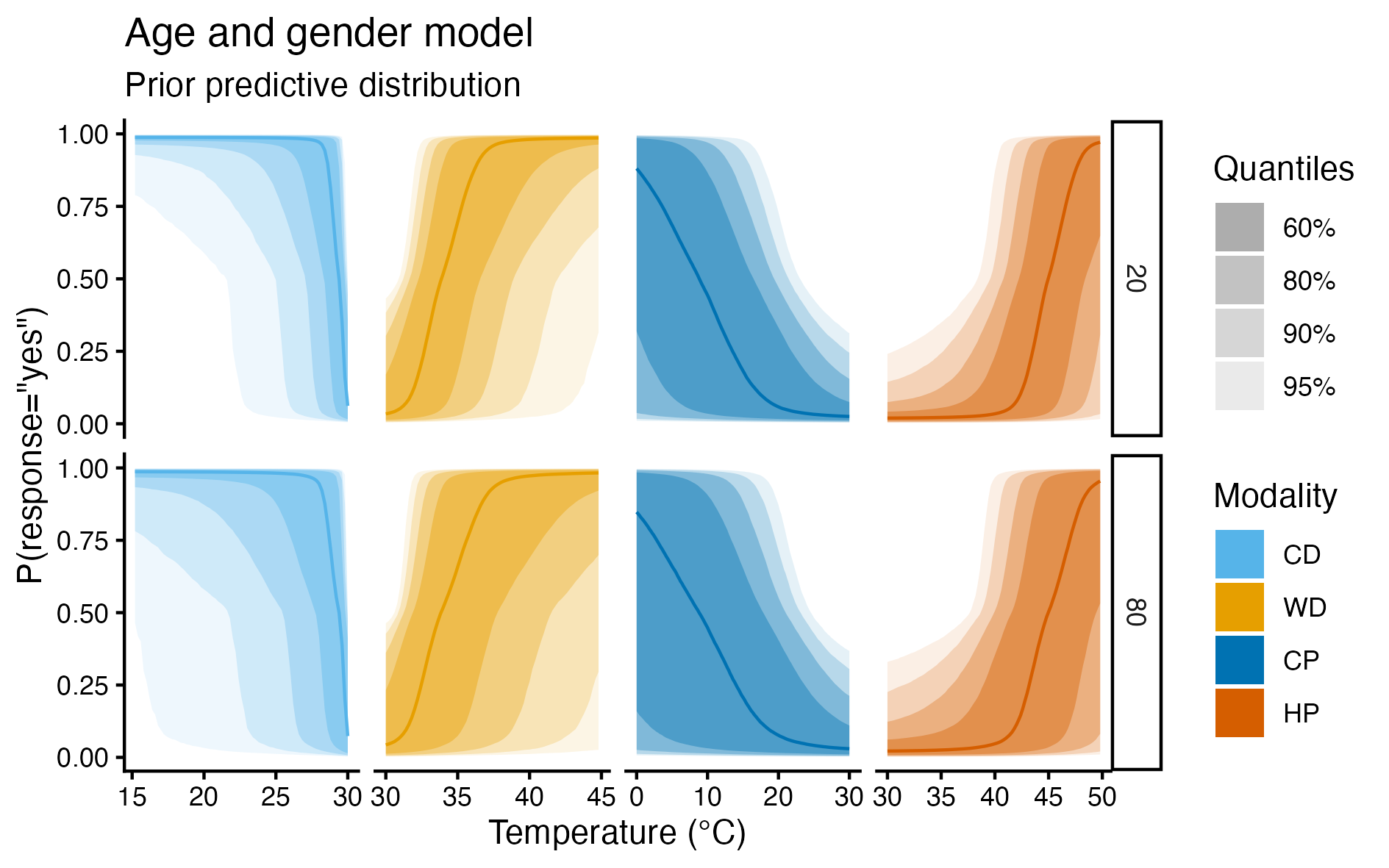

###### ***Figure S1 Prior predictive distribution for the age and gender model, for control participants aged 20 and 80***

### Ageing and neuropathy model

The priors for parameters shared by the ageing only and the ageing and neuropathy model are the same in both models. For the effects of neuropathy (𝜅_s_), we use the same priors as for the effects of gender (𝜅_g_).

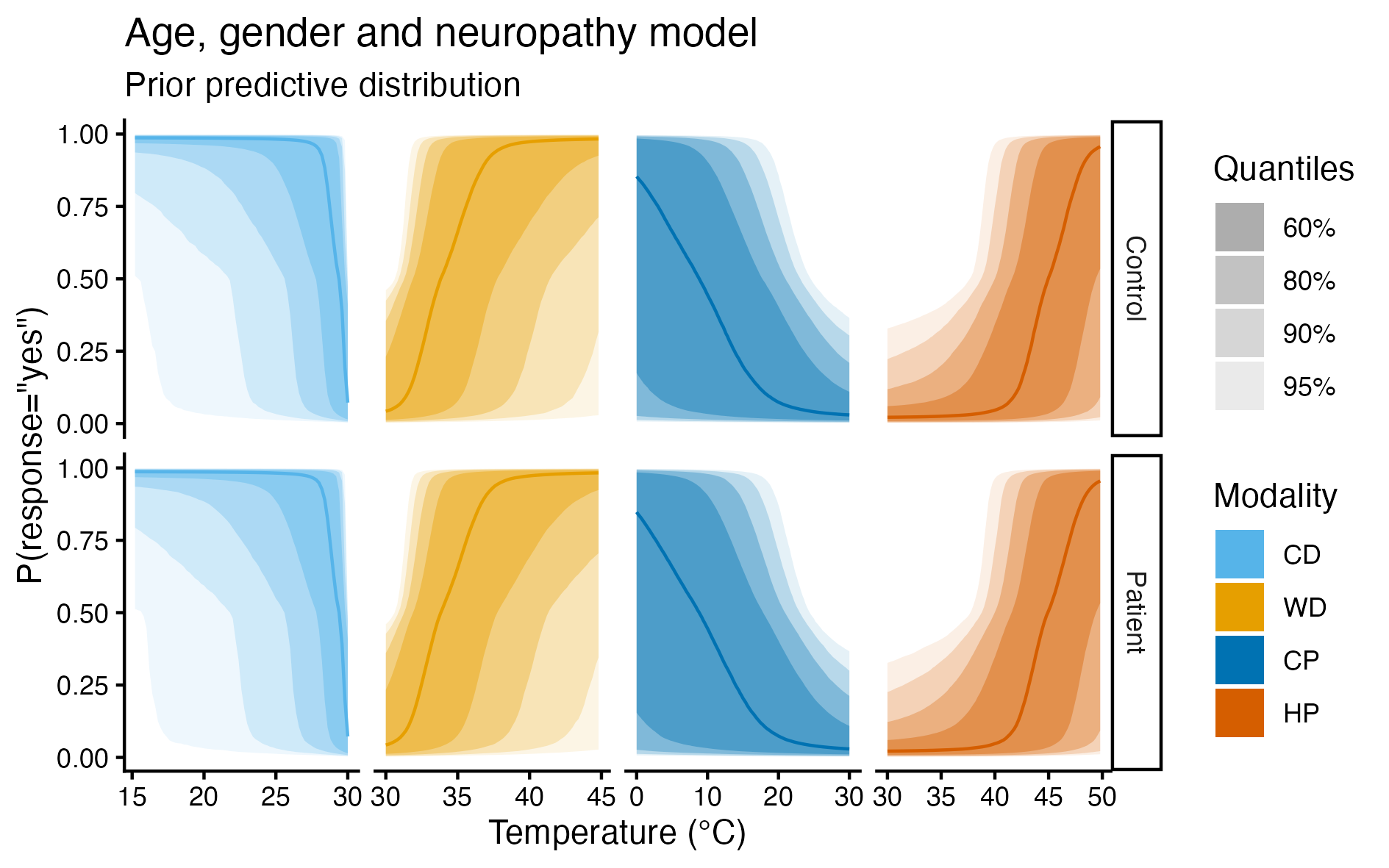

###### ***Figure S2 Prior predictive distribution for the age, gender and neuropathy model - 50 years old participant***

Descriptive statistics on task parameters

***Table S1. Proportions of males and females in the different age groups***

| Group | Total | Males (% group) | Females (% group) |
| --- | --- | --- | --- |
| Controls aged 20 to 35 | 22 | 11 (50%) | 11 (50%) |
| Controls aged 36 to 50 | 16 | 5 (31.2%) | 11 (68.8%) |
| Controls aged 51 to 65 | 14 | 3 (21.4%) | 11 (78.6%) |
| Controls aged 66 to 80 | 23 | 11 (47.8%) | 12 (52.2%) |
| Patients aged 47 to 80 | 33 | 18 (54.5%) | 15 (45.5%) |

***Table S2. Average ramp duration (s) per participant and condition***

| **Task** | **Group Mean** | **SD** | **Minimum** | **Maximum** |
| --- | --- | --- | --- | --- |
| Cold Detection | 1.91 | 1.41 | 0.492 | 8.17 |
| Warm Detection | 2.90 | 1.16 | 0.745 | 6.98 |
| Cold Pain | 8.38 | 2.49 | 1.50 | 11.5 |
| Heat Pain | 5.84 | 1.06 | 2.10 | 7.34 |

***Table S3. Proportion of “yes” responses per participant and condition***

| **Task** | **Group Mean** | **SD** | **Minimum** | **Maximum** |
| --- | --- | --- | --- | --- |
| Cold Detection | 0.666 | 0.137 | 0.333 | 1 |
| Warm Detection | 0.531 | 0.134 | 0 | 0.875 |
| Cold Pain | 0.302 | 0.192 | 0 | 0.575 |
| Heat Pain | 0.416 | 0.101 | 0.05 | 1 |

Effects of gender

### Age and gender model

***Table S4. Effects of gender on psychometric function parameters across modalities***

| **Modality** | **Parameter** | **κ** | **90% CI** | ***P*(𝜅>0)** | **log_10_(BF₀₁)** | **Interpretation** |
| --- | --- | --- | --- | --- | --- | --- |
| Cold Detection | Threshold | 0.045 | [-0.398, 0.481] | .569 | 0.50 | Not different |
|  | Slope | 0.126 | [−0.334, 0.601] | .674 | 0.54 | Not different |
| Warm Detection | Threshold | -0.144 | [-0.395, 0.100] | .164 | 0.34 | Inconclusive |
|  | Slope | −0.108 | [-0.509, 0.283] | .329 | 0.45 | Inconclusive |
| Cold Pain | Threshold | -0.025 | [-0.226, 0.178] | .417 | 0.47 | Inconclusive |
|  | Slope | -0.035 | [−0.309, 0.248] | .410 | 0.49 | Inconclusive |
| Heat Pain | Threshold | -0.008 | [-0.060, 0.043] | .398 | 0.72 | Not different |
|  | Slope | -0.080 | [−0.490, 0.327] | .372 | 0.51 | Not different |

### Age, gender, and neuropathy model

***Table S5. Effects of gender on psychometric function parameters across modalities***

| **Modality** | **Parameter** | **κ** | **90% CI** | ***P*(𝜅>0)** | **log_10_(BF₀₁)** | **Interpretation** |
| --- | --- | --- | --- | --- | --- | --- |
| Cold Detection | Threshold | -0.177 | [-0.541, 0.180] | .210 | 0.48 | Inconclusive |
|  | Slope | 0.274 | [−0.116, 0.669] | .873 | 0.39 | Inconclusive |
| Warm Detection | Threshold | -0.068 | [-0.253, 0.118] | .274 | 0.58 | Not different |
|  | Slope | −0.250 | [-0.595, 0.081] | .110 | 0.25 | Inconclusive |
| Cold Pain | Threshold | -0.053 | [-0.226, 0.114] | .301 | 0.50 | Not different |
|  | Slope | -0.051 | [−0.309, 0.202] | .374 | 0.52 | Not different |
| Heat Pain | Threshold | -0.021 | [-0.065, 0.022] | .217 | 0.65 | Not different |
|  | Slope | -0.139 | [−0.537, 0.221] | .290 | 0.48 | Inconclusive |

Posterior predictive plots for individual fits

To verify that our model fit the data reasonably well, we also created posterior predictive plots, showing the response of the participant against what the model assumes their psychometric functions to be.

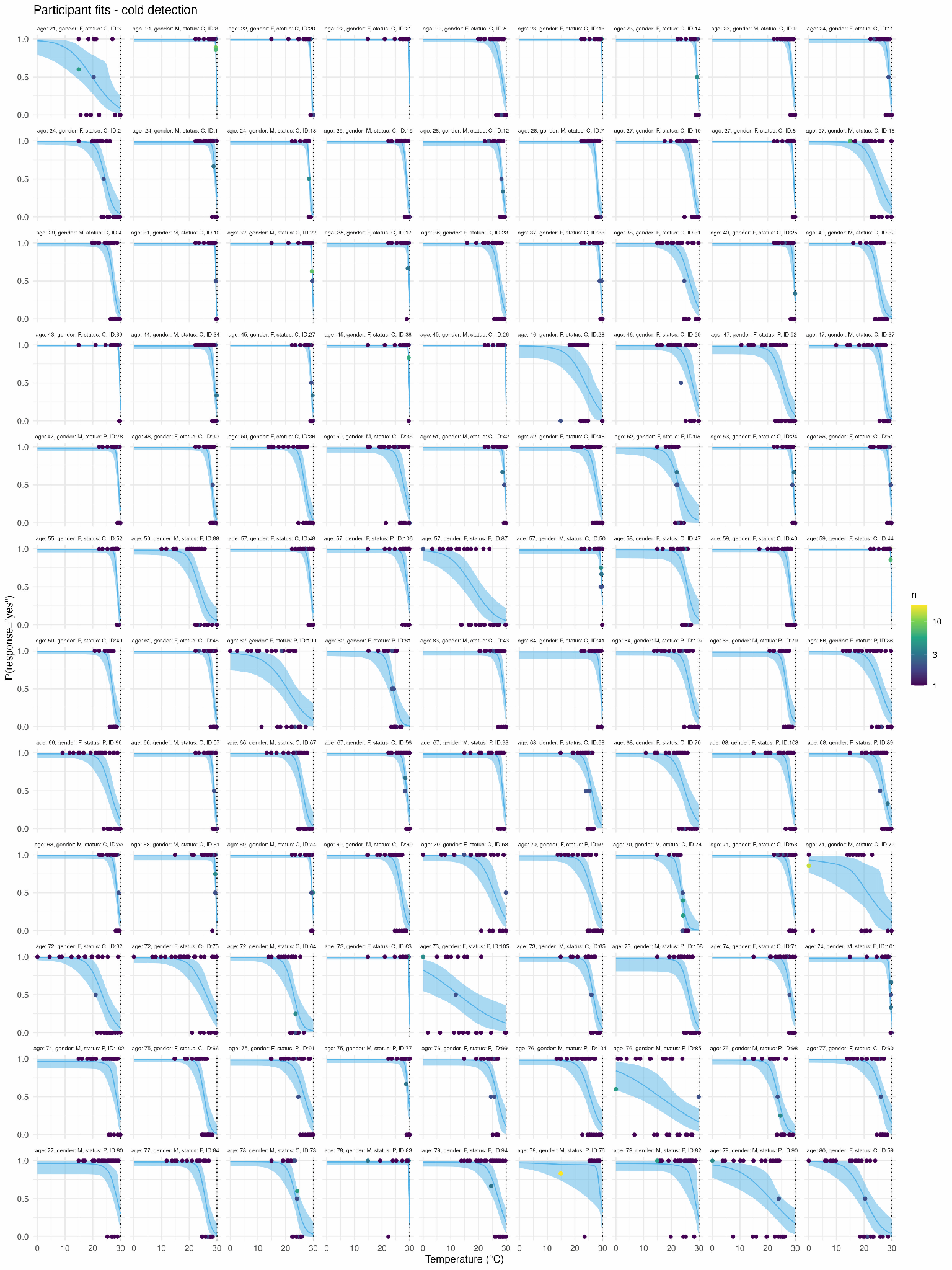

###### ***Figure S3 Posterior predictive plots for cold detection***

*Psychometric function (line: posterior mean, shaded area: 95% CI) and responses (dots) for each participant.*

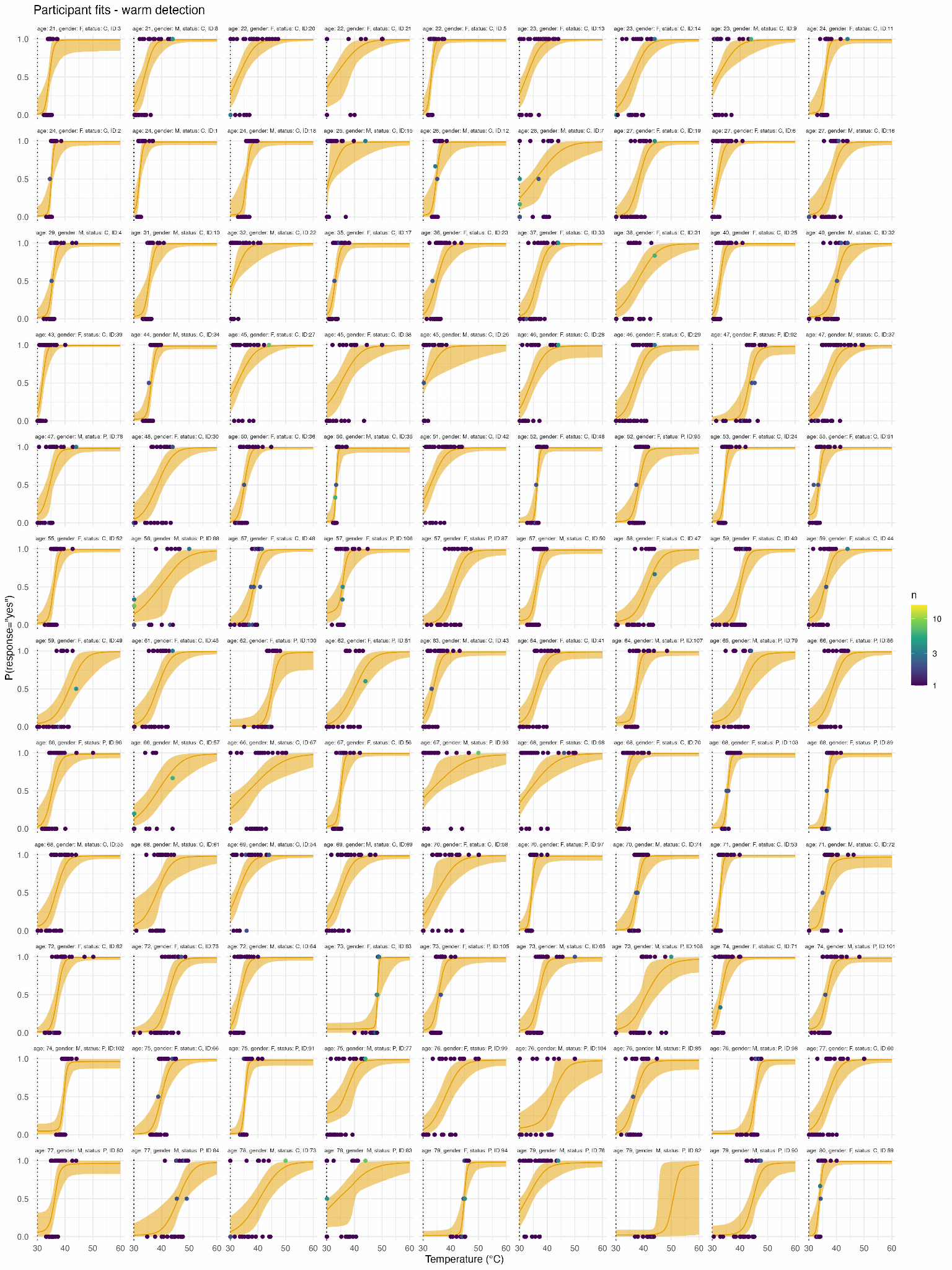

###### ***Figure S4 Posterior predictive plots for warm detection***

*Psychometric function (line: posterior mean, shaded area: 95% CI) and responses (dots) for each participant.*

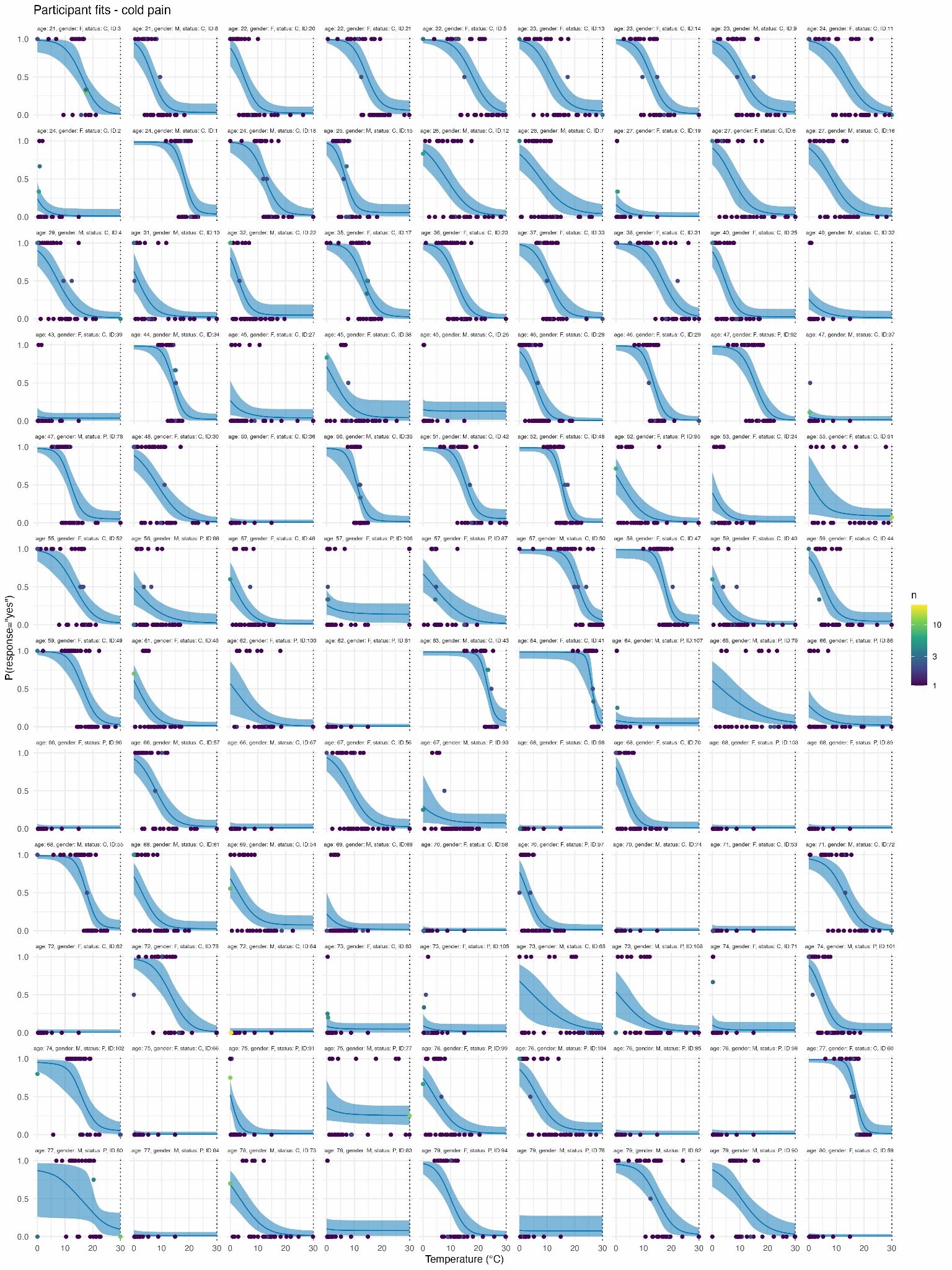

###### ***Figure S5 Posterior predictive plots for cold pain***

*Psychometric function (line: posterior mean, shaded area: 95% CI) and responses (dots) for each participant.*

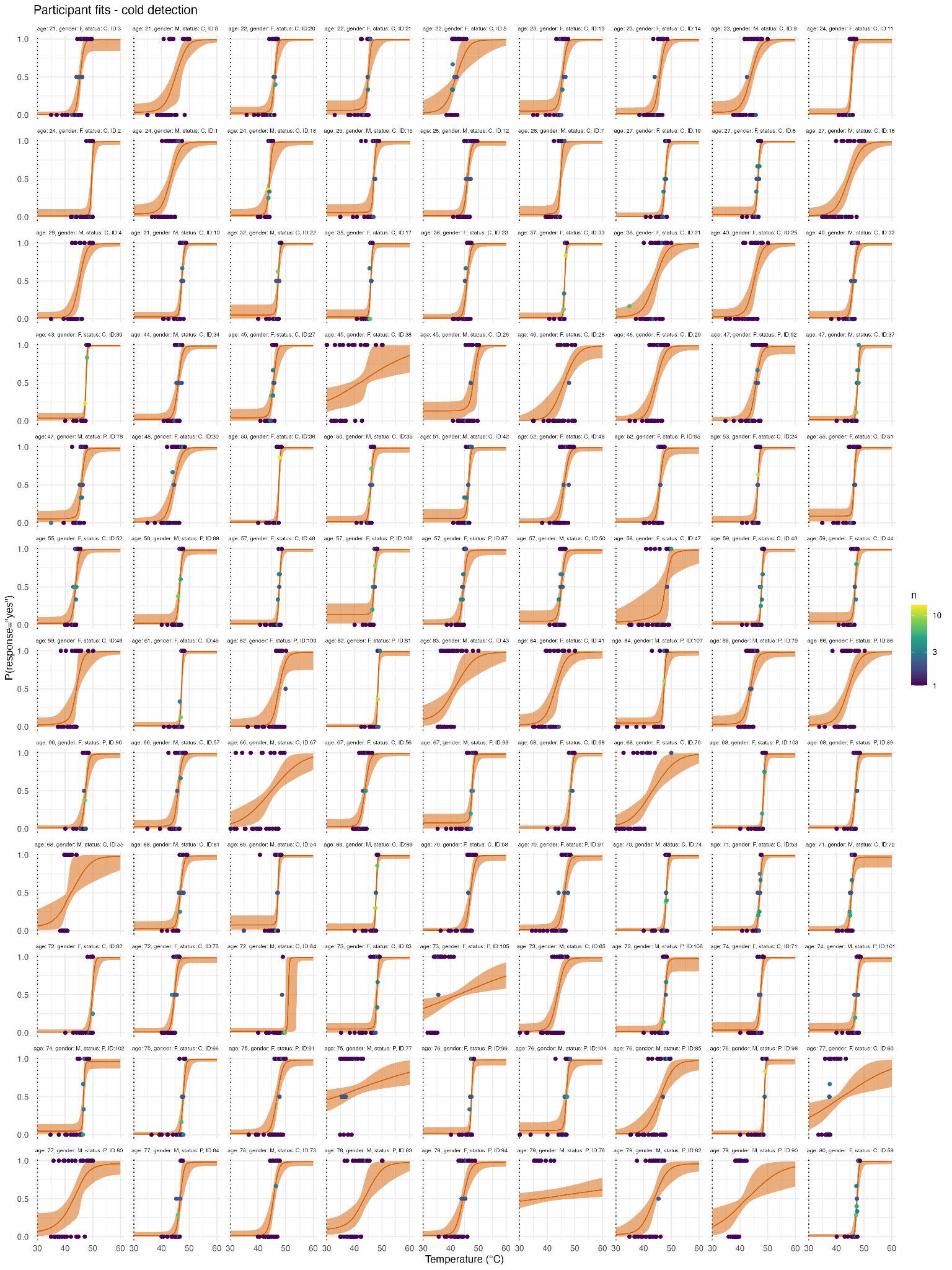

###### ***Figure S6 Posterior predictive plots for heat pain***

*Psychometric function (line: posterior mean, shaded area: 95% CI) and responses (dots) for each participant.*

Receiver Operating Characteristics Curves

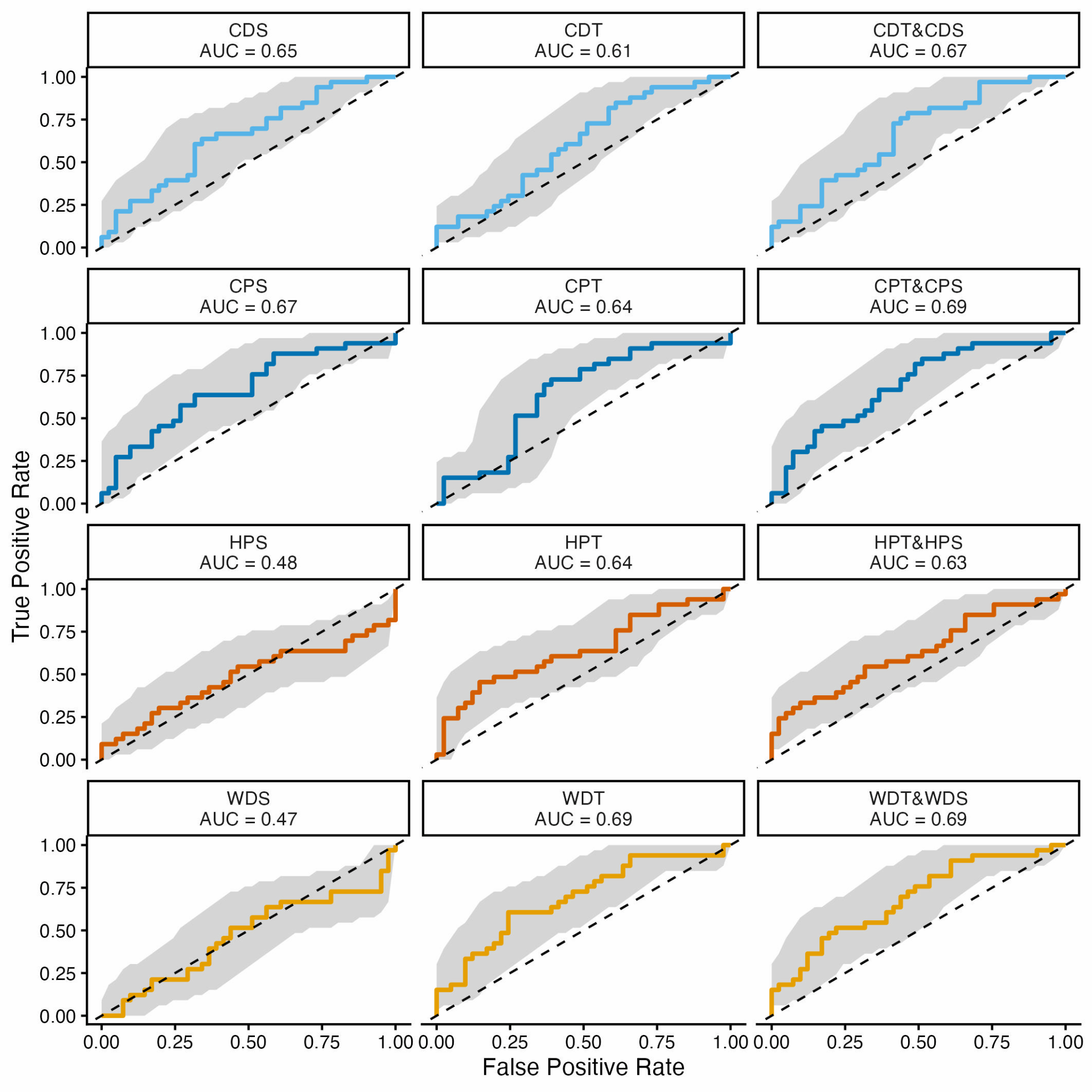

###### ***Figure S7 ROC curves for each of the single task classification models***

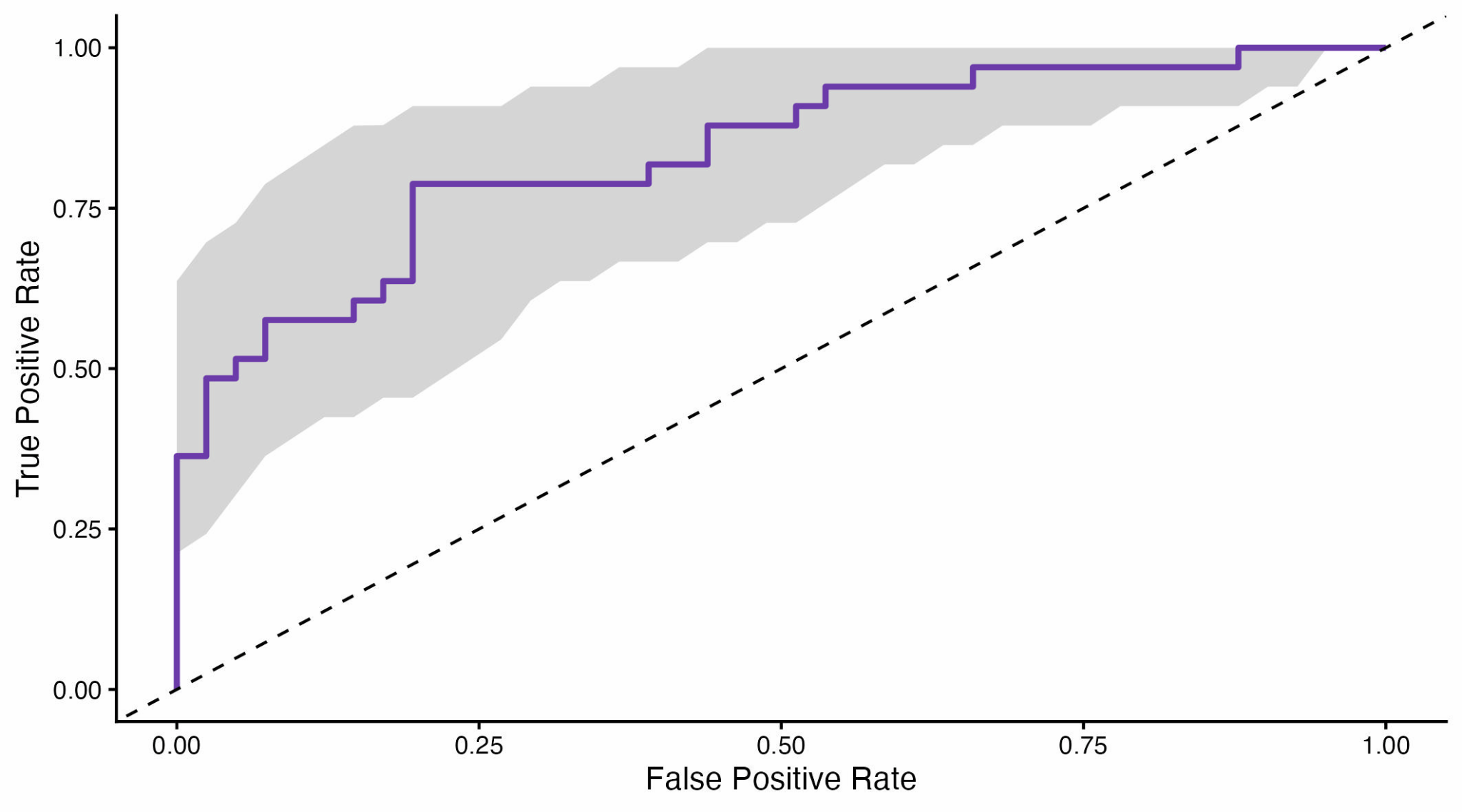

###### ***Figure S8 ROC curves for the all tasks classification models***
